# Engineering Infrared Light Detection in Blind Human Retina Using Ultrasensitive Human TRPV1 Channels

**DOI:** 10.1101/2025.01.08.631889

**Authors:** Morgan Chevalier, Firas Fadel, Tímea Májer, Dániel Péter Magda, Lili Gerendás, Ferenc Kilin, Zoltán Zsolt Nagy, Arnold Szabó, Botond Roska, Guilherme Testa-Silva

**Affiliations:** Institute of Molecular and Clinical Ophthalmology Basel, Basel, Switzerland; Department of Anatomy, Histology and Embryology, Semmelweis University, Budapest, Hungary; Department of Ophthalmology, Semmelweis University, Budapest, Hungary; Department of Ophthalmology, University of Basel, Basel, Switzerland

**Keywords:** TRPV channels, Vision restoration, Human retina, Photothermal stimulation, Protein engineering, Engineered ion channels, Electrophysiology in retinal cells, Gene therapy

## Abstract

Engineering infrared light sensitivity in the blind human retina could restore visual function in patients with regional retinal degeneration. However, current approaches are complex and contain non-human biological components. Using rational protein design we engineered human TRPV1 channels (Δ786-840) with temperature sensitivity shifted from 45 to 41°C that enabled near-infrared light-induced heat activation of mammalian cells at close to physiological temperatures. When expressed in ganglion cells of human retinal explants, Δ786-840 TRPV1 generated robust spiking responses to brief near-infrared light-induced temperature transients. Additionally, increasing intensity of radiation evoked graded responses correlating with increasing firing frequencies. Unlike previous approaches that used non-human TRPV1 channels, which risk immune reactions and a multicomponent system that poses barriers to clinical implementation, this single component human-derived approach eliminates immunogenicity concerns, addressing a major challenge to clinical translation, and allow gene delivery using adeno-associated viral vectors.

## Introduction

Restoring vision in blind individuals with retinal degenerative diseases remains one of the crucial challenges in translational neuroscience and ophthalmology^1^. These diseases often lead to the loss of function or loss of photoreceptors, the primary cells responsible for capturing and converting light into visual percepts^2^. Optogenetic therapies that target light sensitive channels or pumps to retinal cells in blind retinas are promising approaches to restore vision and are in clinical trials^3,4^. However, in several forms of retinal degenerations, such as age related macular degeneration, the loss of light sensitivity is partial, with a mosaic of light-sensitive and light-insensitive areas within the retina^5^. This residual photoreceptor activity presents challenges for optogenetic techniques that rely on external bright visible light stimulation.

One way to address this problem is the targeted expression of thermosensitive ion channels together with heat-stimulation of light insensitive retinal regions using infrared or near-infrared radiation. Indeed, thermosensitive Transient Receptor Potential (TRP) channels are used by several species, such as pit vipers and vampire bats, to capture infrared light to perform visual behaviors, such as foraging, hunting prey, or seeking shelter^6–8^. A recent vision restoration approach^9^ has utilized rat and snake TRP channels for heat-based vision restoration. However, this system uses two components—gold nanorods and non-human TRP channels—which complicates therapeutic delivery, and a non-human protein, which poses a risk for immune reaction.

To address this problem, we aimed to use a human TRP channel, alone. However, the most well-characterized human thermosensor channel, TRPV1, has a very high—~45°C^10,11^— activation threshold as it functions to detect noxious heat stimuli^10,12–14^. To overcome this limitation, we aimed to engineer a human TRPV1 channel (hTRPV1) with a lower temperature threshold suitable for infrared radiation-based vision restoration. By aligning the hTRPV1 sequence with orthologs from multiple species, we identified key differences in the N- and C-termini that modulate temperature sensitivity. We also drew from previous studies detailing the mechanisms underlying TRP thermal responses^14,15^. Building on these insights, we engineered human TRPV1 channels with enhanced temperature sensitivity, lowering their temperature of half activation (T_50_) from around 45°C to 41°C. This reduction brings the activation temperature closer to the safety margin for clinical applications, where controlled thermal stimulation of retinal cells is required without damaging surrounding tissues^16^.

## Results

We aimed to engineer hTRPV1 with increased heat sensitivity. To achieve this, we aligned sequences of hTRPV1 with orthologs from rat (*Rattus norvegicus*) mouse (*Mus musculus*) ground squirrel (*Ictidomys tridecemlineatus*), camel (*Camelus bactrianus*), and vampire bat (*Desmodus rotundus*) (Fig. 1a). The N-terminal positions N125 and E189 (Fig. 1a, upper panel, b, e) were of particular interest, given their reported roles in modulating temperature sensitivity in mouse and ground squirrel TRPV1^17^. Additional positions of interest were the highly conserved residues R772 and R782 (Fig. 1a, lower panel, b, e), which have been implicated in PIP2 binding^15^, a lipid that increases the thermal activation threshold of TRPV1 by negatively regulating TRPV1 activity. We hypothesized that mutating these four residues would decrease the thermal activation threshold in hTRPV1.

**Fig. 1.**
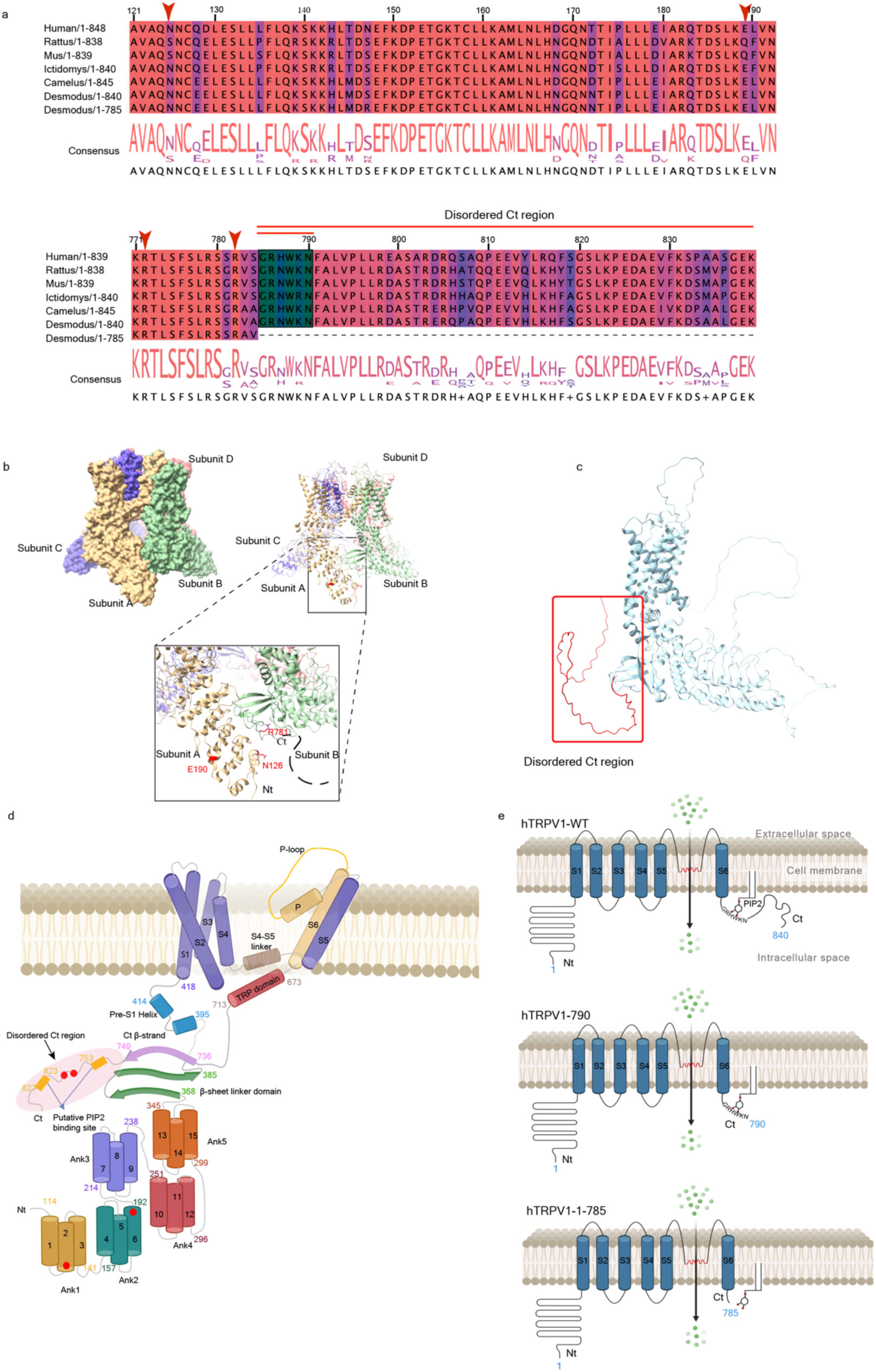
Structural and sequence comparison of TRPV1 channels and synthetic rationale. **a,** Multiple sequence alignment of TRPV1 across six species, including Human, Rattus norvegicus (Rat), Mus musculus (Mouse), Ictidomys tridecemlineatus, Camelus bactrianus, and Desmodus rotundus (both long and short sequences). The upper panel shows the alignment of the N-terminal region (122–193), and the lower panel displays the alignment of the disordered Ct region (780–840). Residues N125, E189, R772, and R782, which have been reported to be implicated in TRPV1 temperature sensitivity, are indicated with red arrows. The disordered Ct region is marked, and the GRxWKN motif is highlighted in green. A consensus sequence is shown below each alignment. **b,** Cryo-EM structure of human TRPV1 (PDB ID: 8GF8): Surface (left panel) and cartoon (right panel) representations of the four subunit arrangements (A, B, C, and D). The inset zooms into the N-terminal region of subunit A and the Ct region of subunit B, with residues N125, E189, and R772 colored in red. The disordered C-terminal region (790–840) is not visible in the structure and is indicated by a black line. **c,** AlphaFold model of a full-length hTRPV1 subunit, highlighting the disordered Ct region (in red). **d,** Schematic representation of TRPV1 structure, showing the transmembrane domains (S1–S6), ankyrin repeats (Ank1– Ank5), the disordered Ct region, and two putative PIP2 binding sites. Location of N125, E189, R772, and R782 residues involved in temperature sensitivity are also indicated with red spheres. **e,** Schematic model showing the interaction between TRPV1 and the PIP2 lipid through the GRxWKN motif. Differences between human TRPV1-wild-type, hTRPV1-790, and hTRPV1-785 variants are shown, highlighting the truncation of the C-terminal tail and its impact on the interaction in each variant.

We first assessed the sensitivity of control hTRPV1 channels to transient heat stimuli in HEK293T cells. To induce rapid thermal transients, we used a custom-designed near-infrared laser system emitting light at 1450 nm. We guided the laser light to cells through a 200 μm core diameter multimode optical fiber positioned ~200 μm from the target cells and confirmed the induced temperature changes at the position of the target cells using a thermal camera. We recorded heat-evoked hTRPV1 channel currents using whole-cell patch-clamp electrophysiology. In all experiments, we had internal controls, HEK293T cells expresing wild-type (WT) hTRPV1, which consistently showed a half-activation temperature (T₅₀) around 45°C. We then compared the properties of wild-type hTRPV1 with the quadruple mutant N125S, E189Q, R772A, R782A. There was no significant change (p>0.05, Mann–Whitney U test) in the thermal activation threshold (half-activation temperature) or in the slope of the activation curve of the quadruple TRPV1 mutant compared to wild-type hTRPV1 (Fig.2f).

We next concentrated on the C-terminal cytoplasmic domain due to its reported role in modulating thermal activation threshold. In vampire bats alternative splicing generates a long (TRPV1-L) and a short (TRPV1-S) isoform^14^. TRPV1-S lacks part of the C-terminal domain and has a ~10°C lower thermal activation threshold (30.5 ± 0.7°C) than TRPV1-L (39.6 ± 0.4°C). Similarly, the C-terminal tail of rat TRPV1 has been shown to interact with membrane lipids^15^, including PIP2, to modulate the channel’s sensitivity to thermal stimuli. Since sequence alignment between hTRPV1 and TRPV1 orthologs across multiple species showed that the C-terminal cytoplasmic domain is highly conserved, we asked if truncating the C-terminal region of hTRPV1 that corresponds to the difference between vampire bat TRPV1-L and TRPV1-S isoforms could affect channel activation threshold or slope. We focused on residues 786 to 840, a region predicted to be disordered and, correspondingly, could not be modeled in the recently published Cryo-EM structure of hTRPV1^18^ (Fig. 1b, c, e). We created the deletion mutant Δ786-840 hTRPV and compared its heat activation threshold to wild-type hTRPV1. Δ786-840 hTRPV had a ~4°C decrease in thermal activation threshold (41.23 ± 1.01°C) relative to wild-type hTRPV1 (45.0 ± 0.7°C)(Fig. 2d). These findings suggest that the disordered C-terminal domain of hTRPV1 contributes to thermal activation regulation. Since the difference between TRPV1-S and TRPV1-L thermal activation thresholds is larger than 4°C in vampire bats, additional structural features likely influence the species-specific tuning of TRPV1 activation thresholds.

**Fig. 2.**
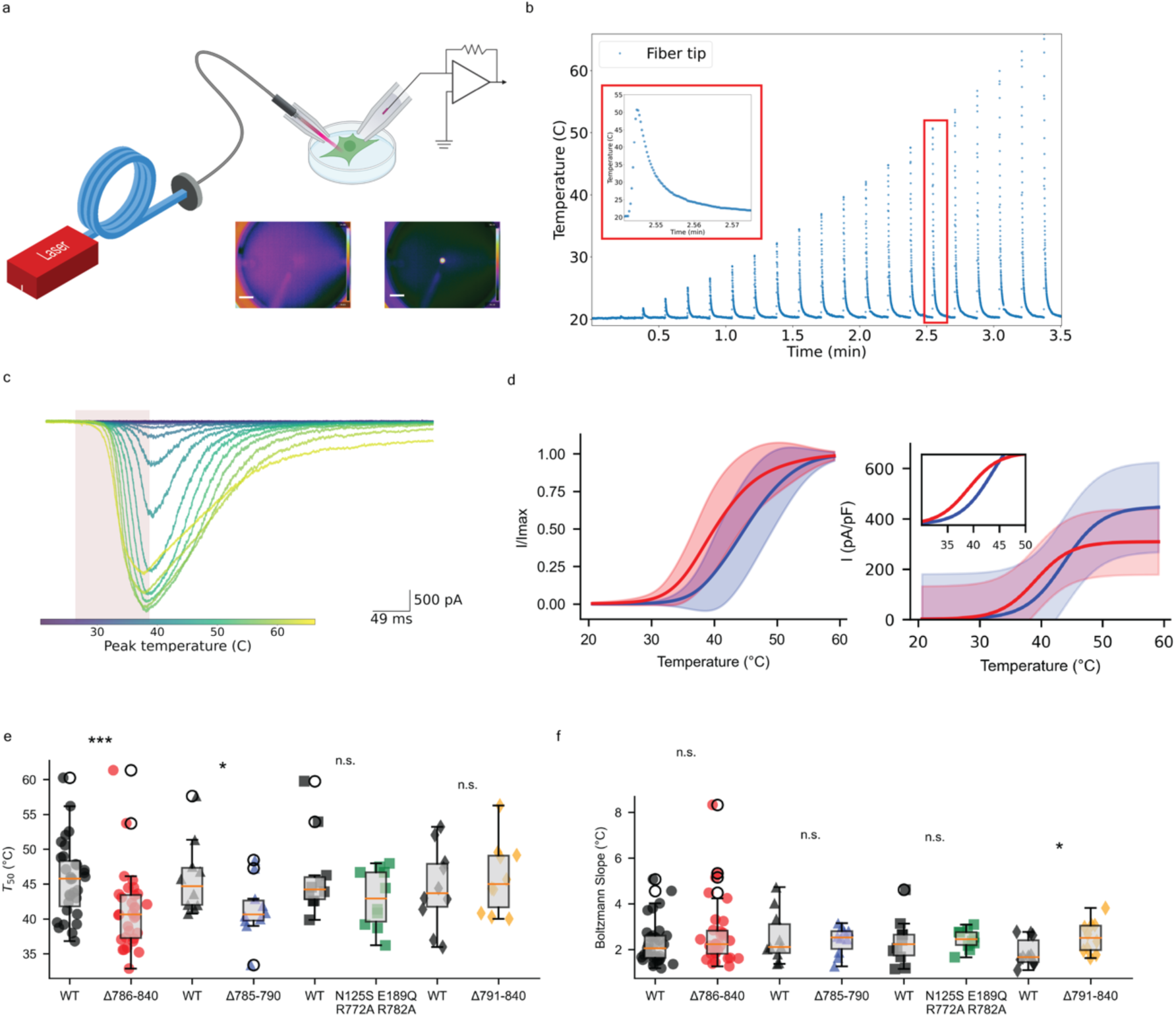
Electrophysiological features of hTRPV1 and mutants on HEK293T cells. **a,** Schematic of an experimental setup used to elicit brief (100 ms) temperature transients on HEK cells via a bare optical fiber coupled to a 1450 nm (10W) laser. Patch-clamp electrodes record the channel response to the heat stimulus. The lower right images (Scale bar: 5 mm) are taken from a FLIR microbolometer thermal camera, showing the temperature distribution before (left) and during (right) laser activation, with the bright spot indicating localized heating at the fiber tip. **b,** Time series of temperature measurements recorded using a FLIR microbolometer thermal camera near the bare end of the optical fiber, where the heat stimulus was delivered to cells. The repeated temperature transients indicate pulsed heat delivery, with maximum values reaching above 60°C at the fiber tip. The inset shows a magnified view of the rapid temperature decay following a heat pulse, illustrating the kinetics of thermal dissipation in the vicinity of the fiber tip. **c,** Current responses recorded from HEK293T cells in voltage-clamp following heat stimuli of increasing magnitude. The shaded region indicates the time window when the heat stimulus was applied. **d,** Heat-evoked current responses from wild-type hTRPV1 (blue) and the Δ786-840 mutant (red), recorded in HEK293 cells. The left plot shows normalized current (I/Imax) as a function of temperature, with Boltzmann. The right plot shows current density (I in pA/pF) versus temperature. The inset magnifies the activation region between 35°C and 50°C. Shaded areas indicate the variability of the data (mean ± SEM), reflecting differences in activation thresholds and current magnitudes between wild-type and mutant channels. **e,** Temperature thresholds for activation extracted from Boltzmann fits for various TRPV1 mutants compared to their paired wild-type controls. Significant differences between mutants and wild-type are indicated, with ***p < 0.001 and *p < 0.05, determined by the Mann-Whitney U test. Mutants Δ786-840 (mean T_50_ = 41.23±1.01°C) and Δ785-790 (T_50_ = 41.39±1.34°C) show statistically significant reductions in threshold temperature compared to wild-type (T_50_ = 45.57±0.7°C). **f,** Maximum velocity (slope). The Δ791-840 mutant shows a significantly increased slope (2.54±0.22°C) compared to wild-type (1.87±0.19°C), indicating a change in the steepness of channel activation kinetics for this specific mutant.

A potential explanation for the decreased activation threshold of the Δ786-840 hTRPV1 mutant is that the 786-840 region, similar to TRPV in rats, mediates interactions with membrane lipid PIP2. The precise location of the PIP2 interaction domain is however unknown^15,19,20^. Through sequence analysis, we identified a conserved positively charged GRxWKN motif corresponding to residues 785-790 in the C-terminal tail of hTRPV1, which could mediate PIP2 binding. We therefore generated two mutants: Δ791-840 hTRPV1 and Δ785-790 hTRPV1 variants. In Δ791- 840, the GRxWKN motif is retained while truncating the rest of the disordered C-terminal domain. Conversely, in Δ785-790, the GRxWKN motif is deleted but the remainder of the C-terminal domain is retained. Electrophysiological analysis revealed that Δ791-840 hTRPV1 had a thermal activation threshold (45.92 ± 1.62 °C) comparable to the (paired control) wild-type channel (44.51 ± 1.82°C), (Fig. 1e), whereas Δ785-790 had a lower thermal activation threshold (41.39 ± 1.34°C) when compared to the wild-type (45.90 ± 1.67°C). The effects observed in both Δ785-790 and Δ786-840 variants underscore the importance of the GRxWKN motif in controling TRPV1 heat sensitivity likely via direct interaction via PIP2.

We focused our attention on a particular mutant Δ786-840 (shown in Figure 2d), which switches on at a mid-point temperature (T_50_) of 41.23 ± 1.01°C. This activation temperature is slightly above the human body’s normal physiological temperature of roughly 37°C, but still below the threshold where retina tissues risk permanent damage, such as blood clotting or protein denaturation^16^. After confirming these properties in HEK293T cells, we moved to experiments using postmortem human retina samples (shown in Figure 3). Our goal was to use a near-infrared (NIR) laser to activate this engineered system, relying on just a single genetic modification and a one-time treatment. In other words, we wanted to see if we could make human retinal tissue sensitive to gentle heating from an NIR laser using only this single-component mutant. The Δ786-840 mutant was chosen because its activation profile suggests it might respond well to moderate increases in temperature. By studying how this mutant behaves in human retina samples, we aimed to determine whether it could be safely and reliably triggered by NIR light without causing irreversible damage. In practical terms, this approach could help us control cellular responses in the retina using small, precise temperature changes, potentially guiding future clinical efforts into restoring or modulating vision in damaged retinal tissue.

**Fig. 3.**
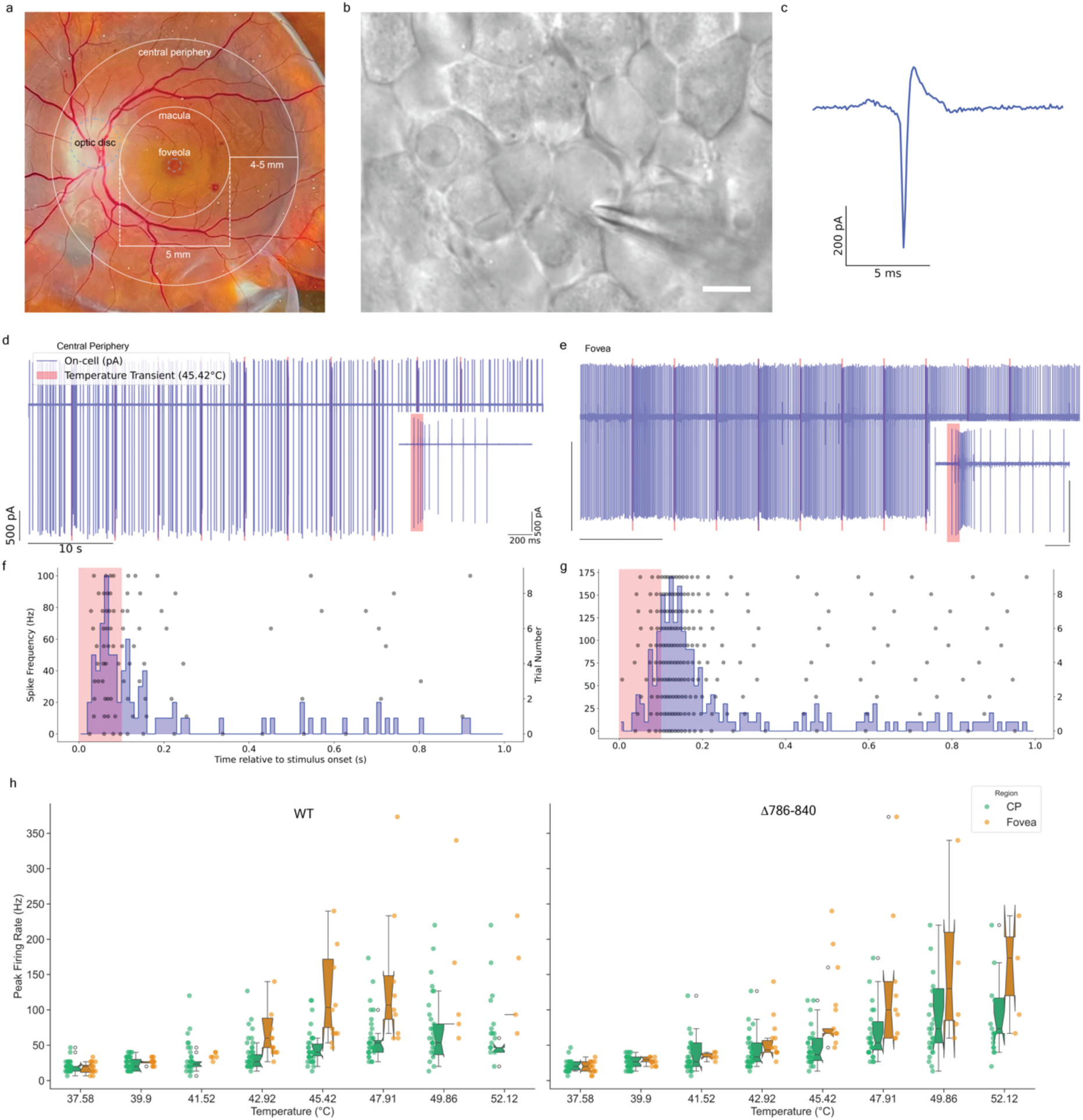
Diversity of Spontaneous and Evoked Action Potentials in Human Retinal Ganglion Cells (*ex vivo*). **a,** Schematic representation of the human retina, highlighting the fovea and central retinal areas from which recordings were obtained. **b,** Differential interference contrast (DIC) image of human retinal ganglion cells, showing a patch pipette positioned for recording. The cells display typical polygonal morphology. Scale bar: 10 µm. **c,** Juxtasomal recording and whole-cell current-clamp recording of a spontaneous action potential from two distinct human retinal ganglion cells, illustrating the fast kinetics of the spike with a narrow half-width. **d,** One-minute juxtasomal recording from a human retinal ganglion cell in the central periphery, showing spontaneous action potentials and in response to heat transients (shaded red rectangles) at 52.12°C. The magnified portion on the right shows the fine temporal resolution of the firing response during and after a temperature transient. **e,** Juxtasomal recording in the macula, heat transients (shaded red rectangles) at 45.42°C. Similar to panel (d), this trace captures the firing pattern during baseline conditions and its modulation by repeated thermal stimuli. Inset highlights a high- frequency burst in response to the heat transient. Scale bar dimensions are consistent with panel (d). **f,** Peri-stimulus time histogram (PSTH) and raster plot of spike frequency in response to heat transients from the retinal ganglion cell recording in panel (d). Individual trials are represented by gray dots in the raster plot overlay, showing variability in spike timing across trials. **g,** PSTH, and raster plot of spike frequency in response to heat transients from the retinal ganglion cell recording in panel (e). **h,** Peak firing rate of retinal ganglion cells in response to increasing temperature transients, comparing wild-type and Δ786-840 mutant hTRPV1 channels. Recordings from cells in the central periphery (CP, green) and macular(orange) regions. The notched box plots show the distribution of peak firing rates at different stimulus temperatures, with larger firing rates observed in the macula, particularly at higher temperatures. Despite a trend for macular RGCs to exhibit higher frequency bursts, no statistically significant differences were found between groups. See Supplementary Table 1 for detailed statistical analysis.

Therefore, we transduced hTRPV1 into adult human retinal explants cultured ex vivo for up to four weeks after postmortem extraction. As previously documented, human retinal explants lose normal light-evoked responsiveness within few days of isolation in culture media^21^. Using AAV delivery with the human synapsin promoter, we transduced human retinal ganglion cells (RGCs) with either the wild-type hTRPV1 ortholog or our selected mutant Δ786-840. To measure heat-driven (i.e., NIR-driven) responses in the human retina while minimizing the impact of the recording apparatus, we performed juxtasomal recordings from visually identified RGCs (Fig. 3b,c). We applied transient heat pulses in a manner similar to the custom experimental setup previously used on HEK cells. By consistently positioning the bare fiber tip 200μm away from the soma of the target cells, we were able to evoke heat-driven spikes and characterize their electrophysiological properties as a function of increasing temperature transients.

While measuring different RGC subtypes along the macula-to-periphery axis, we observed a variety of response profiles (Fig. 3d,e). Initially, we attributed this diversity to region-specific cell types. Although macula RGCs appeared to exhibit more high-frequency bursts (Fig. 3h), statistical analysis showed no significant differences between regions (Supplementary Table 1). We reliably evoked heat-driven responses in naïve human RGCs due to the presence of native TRP channels. Using a 100 ms heat stimulus repeated across ten trials within one minute at each temperature (Fig. 3d–g), we characterized the properties of the wild-type and Δ786–840 constructs, comparing the mutant channel’s ability to enhance cell excitability in response to heat transients.

Finally, we pooled data from our recordings of human RGCs in both the macula and central periphery (Fig. 4) to compare activation thresholds and response dynamic range, from a rate-coding perspective, between the two constructs (wild-type and Δ786–840) and the untreated postmortem human retina. Our findings confirmed that the untreated retina is mildly heat-sensitive, which is not surprising given that several channels from the TRP superfamily are natively expressed in the human retina. When comparing heat-driven responses among the three groups (Fig. 4a), we observed that at 41.52 °C, the Δ786–840 mutant began to exhibit a more consistent activation pattern in response to heat transients. Next, to test this hypothesis, we calculated z-scores of firing rates (Fig. 4b) by normalizing each cell’s response to its own baseline activity measured just before the stimulus onset (within a 50 ms window). This method accounts for variability in baseline firing rates and fluctuations among different cells and trials. Expressing firing rate changes in terms of standard deviations from the baseline mean provides a standardized metric that makes responses comparable across different cells and experimental groups (wild-type, Mutant, and Control). The departure from the baseline observed in the raster plots was confirmed in the z-score comparison (Fig. 4b), providing a more accurate comparison of spiking responses between groups and practically isolating the effect of the heat stimulus from other sources of variability. Additionally, we measured the peak firing rates during the interval following the heat stimulus onset and plotted the results for all groups (Fig. 4c). As expected, at 41.52 °C, the Δ786–840 mutant displayed a higher mean peak firing rate (32.87 ± 4.28 Hz) compared to the wild-type (18.80 ± 2.60 Hz).

**Fig. 4.**
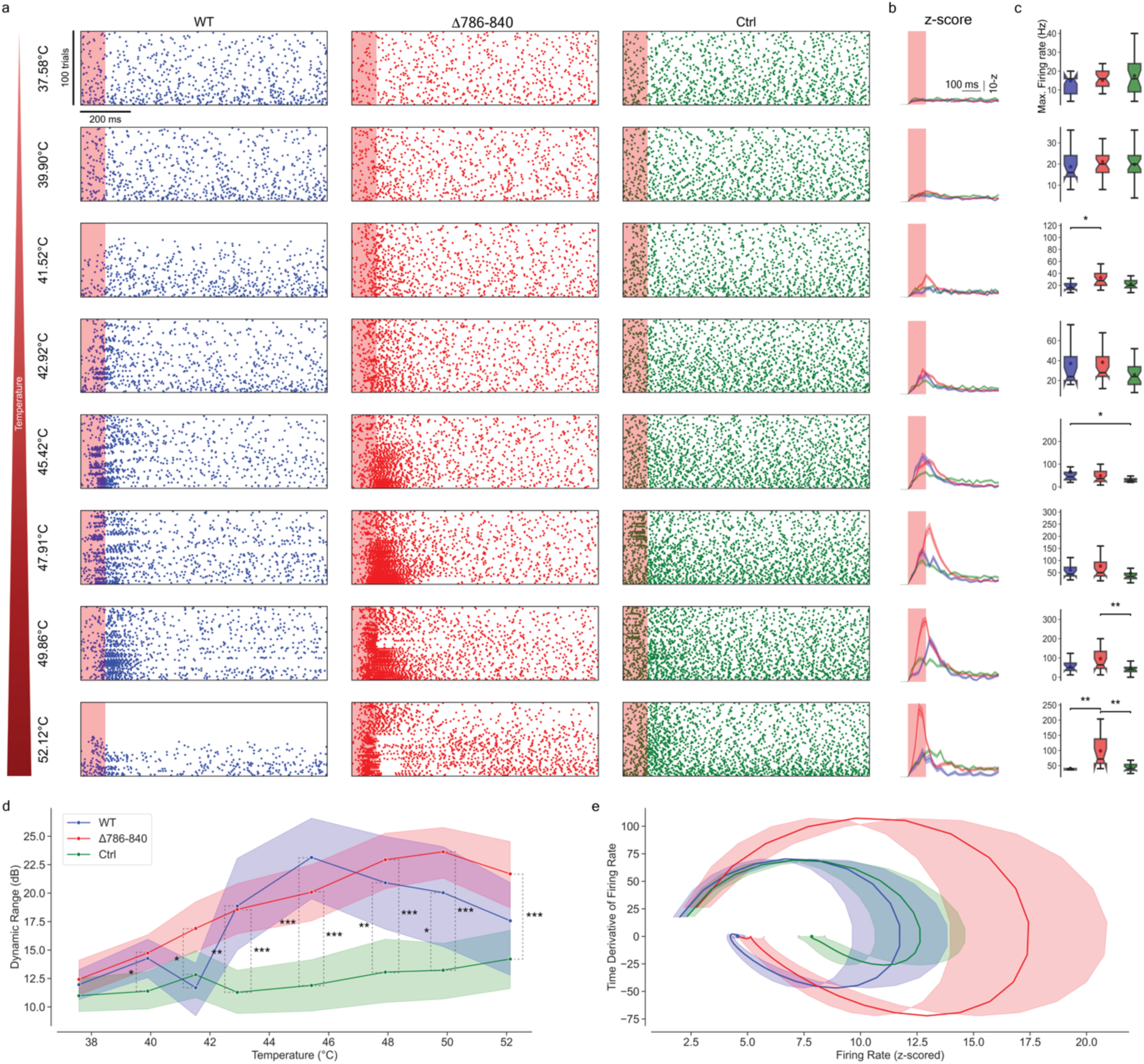
Δ786-840 TRPV1 Mutation Confers Lower Activation Thresholds and Increases Temperature Responsiveness in Human Retina. **a,** Raster plots of pooled recordings from human retinal ganglion cells (RGCs) in the macula and central periphery (CP) regions. The leftmost bar indicates the increasing temperature stimuli applied in each row, ranging from 37.58°C to 52.12°C. Columns represent responses from cells expressing wild-type hTRPV1 (blue), Δ786-840 mutant (red), and control cells without transduction (green). The Δ786-840 mutant shows robust and sustained spiking activity at lower temperatures compared to wild-type hTRPV1 and control cells. The heat transient period is indicated by the red-shaded region. **b,** Z-scored firing rates of RGCs in response to increasing temperature stimuli, shown across rows. A higher positive z-score indicates a significant increase in firing rate compared to baseline (0 to 50ms prior stimuli), while a z-score near zero or negative suggests activity is at or below baseline levels. Notably, at 41.52°C, the Δ786-840 mutant shows a pronounced peak in z-score, indicating a lower activation threshold compared to the wild-type and Control groups, which do not show significant increases in activity at this temperature. **c,** Maximum firing rates (Hz) of RGCs in response to increasing temperature stimuli. Statistical comparisons were made using the Kruskal-Wallis test to assess differences in peak firing rates between wild-type, Mutant, and Control groups at each temperature. When significant differences were found (p < 0.05), pairwise comparisons were conducted using Tukey’s HSD post-hoc test, with significance indicated by *p < 0.05, **p < 0.01, and ***p < 0.001. The Δ786- 840 mutant shows higher firing rates at elevated temperatures compared to wild-type and control groups, particularly at 41.52°C, 49.86°C, and 52.12°C. The control group consistently exhibits lower firing rates across all temperatures. **d,** Dynamic range of retinal ganglion cell firing rates as a function of increasing temperature. Dynamic range was calculated as the logarithmic ratio (in dB) between the peak and baseline firing rates. The Δ786-840 mutant exhibits a significantly broader dynamic range. Statistical analysis was performed using the Kruskal-Wallis test to identify overall differences between groups at each temperature, followed by post-hoc pairwise comparisons using Tukey’s HSD test. Benjamini-Hochberg False Discovery Rate (FDR) correction was applied to adjust p-values for multiple comparisons. **e,** Phase plane plot showing the relationship between z-scored firing rates (x-axis) and their time derivatives (y-axis). RGC response evolution over time following temperature stimuli, with the time derivative capturing the rate of change in firing rate. The Δ786- 840 mutant exhibits higher firing rates and faster changes in response to stimuli, as indicated by the larger trajectories. Additionally, the distinct shapes of the trajectories suggest differences in response patterns, with the Δ786-840 mutant showing a sharper, more transient response compared to wild-type and control groups. The steep downward curves in time derivatives suggest faster recovery from peak firing. The broader x-axis spread of the Δ786-840 mutant indicates increased sensitivity, reaching higher firing rates than wild-type and control.

After establishing that an increase in stimulus intensity led to a higher firing rate, we aimed to indirectly test whether this could convey more information about the stimulus to downstream neurons in the visual pathway by measuring changes in dynamic range (Fig. 4d) of human RGCs across the three experimental groups. Dynamic range was calculated as 20×log10 (Peak Firing Rate/Baseline Firing Rate), expressing the ratio in decibels (dB) to reflect the logarithmic nature of neural response sensitivity. This calculation normalizes the RGC response by accounting for its baseline activity, allowing comparison across neurons with varying baseline firing rates. Higher dynamic range values indicate higher variations in neuronal sensitivity to the stimulus. The Δ786-840 mutant shows a significantly higher dynamic range compared to wild-type and control, particularly between 42°C and 50°C, indicating enhanced thermal sensitivity. Statistically significant differences are marked, highlighting the Δ786-840 mutant’s responsiveness to heat stimuli. From a dynamics perspective, examining the phase plot of z-scored firing rates (Fig. 4e), Δ786-840 mutant exhibits greater dynamic fluctuations in firing rate over time compared to wild-type and control groups. This suggests that the mutant has enhanced responsiveness and adaptability to stimuli. Finally, the pronounced loop in the Δ786-840 mutant reflects its increased sensitivity and faster transitions between different firing states, which could be advantageous in applications requiring rapid and precise neural activation, essential in sensory perception.

## Discussion

We developed a novel approach to enhance the thermal sensitivity of human retinal ganglion cells (RGCs) by engineering TRPV1 ion channels with lower activation thresholds. Our Δ786–840 TRPV1 mutant demonstrated a significant shift in activation temperature, reducing the thermal activation mid-point from the wild-type’s ~45 °C to approximately 41 °C. This modification brings the activation temperature closer to a safe range for therapeutic applications^16,22^, allowing effective stimulation without risking thermal damage to retinal tissues or protein denaturation. By expressing these engineered channels in postmortem human retinal explants, we showed that brief, controlled heat transients delivered via near-infrared (NIR) light can evoke robust spiking responses in RGCs. The Δ786–840 mutant exhibited not only increased sensitivity to thermal stimuli but also a greater dynamic range in firing rates compared to the wild-type and control groups. Its favorable biophysical properties— which allow for precise thermal control and prevent collateral damage to retinal tissues—combined with the reduced risk of immunogenicity from using re-engineered human channels less prone to eliciting immune reactions, make it a promising candidate for future gene therapy-based interventions, particularly in vision restoration.

Our findings align with previous research highlighting the potential of TRPV1 channels in sensory modulation^9^. However, we extend this knowledge by demonstrating the feasibility of using an engineered human variant for vision restoration. The use of NIR light offers advantages over visible light-based therapies, including deeper tissue penetration and reduced interference with residual photoreceptor function. Additionally, our single-component system eliminates the need for auxiliary elements such as gold nanorods, simplifying the therapeutic strategy and potentially improving its safety profile.

Moreover, Off-target effects, a common concern in gene therapy, are unlikely in our system for several reasons. The use of adeno-associated viral (AAV) vectors for retinal delivery ensures localized transduction, with minimal systemic exposure due to direct intravitreal or subretinal injection. Additionally, the synapsin promoter restricts gene expression to retinal ganglion cells, further reducing non-specific expression in other cell types. While the untreated retina exhibits mild sensitivity to heat due to native TRP channels, this response is insufficient for therapeutic stimulation. In contrast, the Δ786-840 mutant significantly enhances responsiveness, allowing precise spatial and temporal control over cellular activation as the Cells outside the NIR exposure field remain unaffected. The ability to induce neural activity at lower temperatures minimizes the risk of thermal damage and expands the therapeutic window. Additionally, the observed consistency in activation patterns across different RGC subtypes and retinal regions suggests broad applicability, regardless of cellular heterogeneity within the retina. Together, these factors demonstrate that our human-derived system is well-suited for clinical translation with minimal off-target risks.

From a broader perspective, our work demonstrates the potential of rational protein engineering to modify human ion channel properties for therapeutic purposes, opening new avenues for treating sensory perception deficits. By harnessing and refining the intrinsic capabilities of human proteins, we can develop targeted interventions that closely resemble physiological conditions, potentially reducing adverse effects. This strategy may extend beyond vision restoration, offering insights into modulating neural activity in various central and peripheral neurological disorders where accurate control over single-cell excitability is desired and visible light activation might be undesirable.

## Methods

### DNA Constructs and Viral Vectors

Polynucleotide sequences encoding human TRPV1-wild-type, the Δ786-840 mutant, and the quadruple mutant (N125S, E189Q, R772A, R782A) were synthesized (GenScript, Piscataway, NJ), fused with P2A and GFP tags, and inserted into the pcDNA3.1 vector under the control of the cytomegalovirus (CMV) promoter. The Δ785-790 mutant, also tagged with P2A and GFP, was cloned into pcDNA3.1 using Ligase Independent Cloning (LIC). For viral vector generation, hTRPV1-P2A-GFP and Δ786-840-P2A-GFP constructs were further subcloned via LIC into recombinant adeno-associated virus (rAAV) vector cassettes under the control of the human synapsin 1 promoter. Viral vectors were packaged into the PHP.eB serotype. Primers used for cloning and mutagenesis experiments are listed in the Table below:

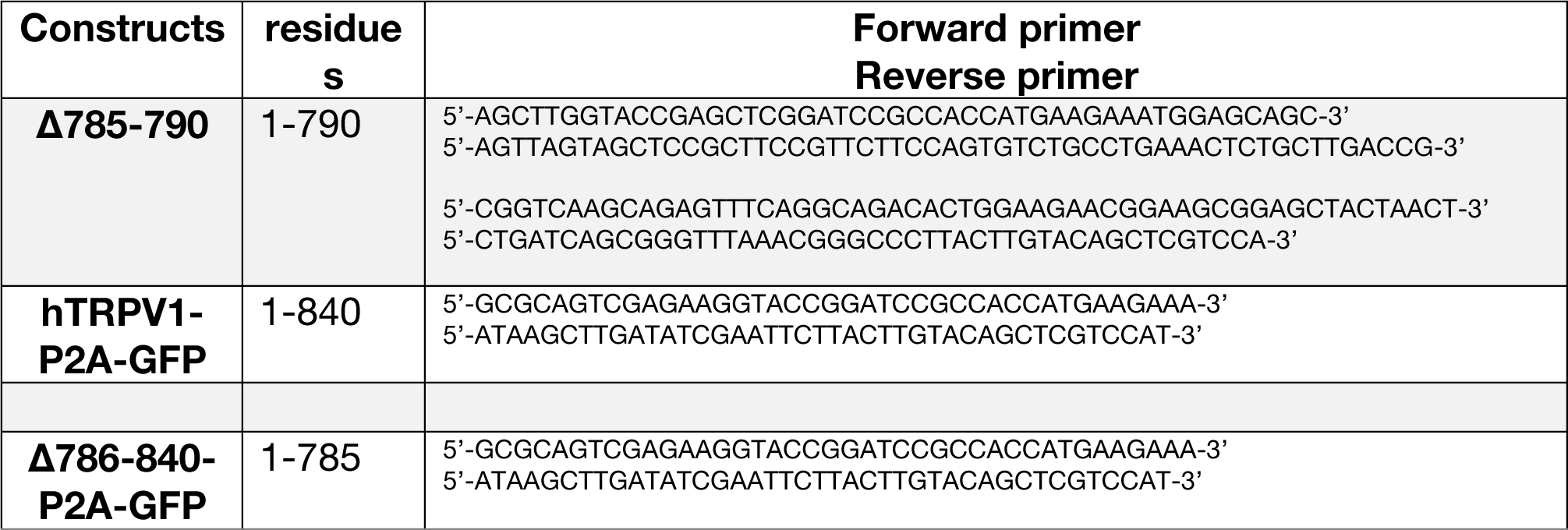

### Bioinformatics

Sequence of *human* TRPV1 (Q8NER1), *Rattus norvegicus* (O35433), *Mus musculus* (Q704Y3), *Ictidomys tridecemlineatus* (I3LZN5), *Camelus bactrianus (*A0A9W3GPJ1), and D*esmodus rotundus*^14^ where aligned using MUSCLE as implemented in Jalview. hTRPV1 model was generated via Alphafold2. Molecular graphics figures were prepared with UCSF ChimeraX (Goddard et al., 2018), and Pymol.

### Cell culture and ion channel expression

HEK293T (ATCC, #CRL-3216) cells were maintained at 37°C and 5% CO_2_ in a humidified incubator. Cells were cultured in DMEM (Gibco, #10566016) supplemented with 10% fetal bovine serum and 1% Penicillin-Streptomycin (Gibco, #15140-122). Cells were passaged after reaching 80% confluency. Cell dissociation was done using 0.05% Trypsin/EDTA (Gibco, # 25300054). For electrophysiological experiments, cells were used from passage 3 to 20.

### Plasmid DNA transfection

For the transient expression of the TRPV1 variants, HEK293T cells were grown in a 35mm culture dish and transfected at 70% confluency with 2.5 µg of plasmid DNA using TransIT-LT1 reagent (Mirusbio, # MIR 2305). Transfection was performed according to manufacturer’s instructions. 24h after transfection, cells were split in 35mm dishes to ensure the isolation of single cells for the electrophysiological recording.

### Human Retinal Tissue

Retinal tissues were obtained from five adult multi-organ donors aged between 43- 63 years, with no documented history of ocular disease. Ethical approval for this study was granted by the Hungarian Scientific and Research Ethics Committee of the Health Sciences Council (ETT TUKEB-34851-2/2018/EKU ETT TUKEB 34851-2/2018/EKU and ETT TUKEB IV/5645-1/2021/EKU) all samples were irreversibly anonymized and processed anonymously to maintain confidentiality, and relevant donor information is summarized in Supplementary Table 2, in accordance with ethical standards. Enucleations were performed by a medical doctor prior to cardiac arrest for the purpose of corneal transplantation. After enucleation, the cornea was transported to the Department of Ophthalmology at Semmelweis University. Concurrently, the posterior segment, including the retina, was transported in Dulbecco’s Modified Eagle Medium/Nutrient Mixture F-12 (DMEM/F-12) to the Retina Laboratory of Semmelweis University. Transportation was conducted under sterile conditions to ensure tissue viability and prevent contamination. Retinal samples were carefully dissected from the macular region, defined as a circular area with an approximate diameter of 5 mm, and the adjacent central peripheral region, which comprises a rim approximately 5–10 mm wide encircling the macula (Fig. 3a). For organotypic retinal culture, 4×4 mm retinal pieces were meticulously isolated to preserve structural integrity and placed photoreceptor-side down on polycarbonate membrane inserts (Corning, 3412). Cultures were maintained at 37 °C and 5% CO₂ in DMEM/F-12 medium (Thermo Fisher Scientific), supplemented with 0.1% bovine serum albumin (BSA), 10 μM O-acetyl-L-carnitine hydrochloride, 1 mM fumaric acid, 0.5 mM galactose, 1 mM glucose, 0.5 mM glycine, 10 mM HEPES, 0.05 mM mannose, 13 mM sodium bicarbonate, 3 mM taurine, 0.1 mM putrescine dihydrochloride, 0.35 μM retinol, 0.3 μM retinyl acetate, 0.2 μM (+)-α-tocopherol, 0.5 mM ascorbic acid, 0.05 μM sodium selenite, 0.02 μM hydrocortisone, 0.02 μM progesterone, 1 μM insulin, 0.003 μM 3,3′,5-triiodo-L-thyronine, 2,000 U penicillin, and 2 mg streptomycin (Sigma-Aldrich). For adeno-associated virus (AAV) infection, 20 μL of viral solution (average titer 10^13^ GC/mL) was applied to each retinal explant one day after plating. The culture medium was renewed every 48 hours. Retinal ganglion cell (RGC) responses were recorded four weeks after virus transduction. The retinal explants were maintained *in vitro* for at least 28 days post-transduction to ensure sufficient expression of the viral construct.

### Electrophysiology in HEK293T

Electrophysiological recordings in HEK293T cells were performed using the patch-clamp technique under whole-cell configuration using an Axopatch™ 200B amplifier (Axon Instruments). Patch pipettes (WPI, # 2BF150-86-10) with resistances of 4 to 7 MΩ were used. Currents were digitized at 20kHz with a low-pass Bessel filter of 10kHz. Data acquisition was made with pClamp 11 (Axon Instruments).

Cells were perfused at room temperature (21±1°C) with an extracellular solution containing (in mM): NaCl 135, KCl 2.5, CaCl_2_ 1, MgCl_2_ 1.6, HEPES 10, D-Glucose 10.

Patch pipettes were filled with an intracellular solution containing (in mM): CsMeSo_4_ 130, CsCl 20, NaCl 9, EGTA 0.2, ATP-Mg 1, HEPES 10.

Heat transients were generated as described below. Light pulses of 100ms with increasing intensities were used to generate heat transients from 20.6 to 59.1°C with a 10s interval. Currents were measured at a holding potential of −60mV.

The maximum current amplitude during each heat pulse was analyzed with Clampfit. The current/temperature curves were fitted to a Boltzmann equation:

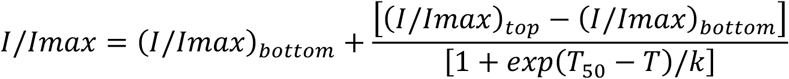

here *I/I* _max_ is the relative current, T_50_ is the temperature to give a half-value of the maximum relative current, and *k* is the slope factor.

### Electrophysiology in Human Retinal Ganglion Cells

Standard juxtasomal patch-clamp recording techniques were employed in these experiments. A Model 700B amplifier (Axon Instruments, Foster City, CA) was utilized, the amplified signals were low-pass filtered at 10 kHz using the built-in 8-pole Bessel filter and sampled at rates between 10 and 50 kHz with a multifunctional data acquisition card (either from National Instruments, Austin, TX, or a Digidata 1440A). Data acquisition was managed using either pCLAMP 10.2 software (Axon Instruments) or custom-developed software capable of synchronous input/output operations and simultaneous control of both the laser and the patch-clamp amplifier. Pipette series resistance and capacitance were compensated using the amplifiers’ internal circuitry, and liquid junction potentials were left uncorrected. Recording pipettes were fabricated from borosilicate glass (Narishige) and had resistances ranging from 1 to 3 MΩ. Currents were typically monitored while clamping at 0 V during juxtasomal recordings. Ames Medium supplemented with L-glutamine was used as the extracellular solution. The same solution was used for both the pipette and the bath. All chemical reagents were purchased from Sigma-Aldrich (St. Louis, MO).

### Fast heat transient generation^17^

To induce transient heat stimuli, a custom-designed laser system emitting at 1450 nm was utilized. The laser diode was driven by pulsed analog modulation, featuring a controller with a negligible rise time to ensure rapid onset of thermal changes. Temperature fluctuations generated by the laser were confirmed using a FLIR A655sc thermal camera equipped with a microbolometer sensor. The laser output was collimated and introduced into a multimode optical fiber with a core diameter of either 50 μm or 200 μm. The distal end of the fiber was stripped and cleaved to expose the fiber core directly. This fiber was mounted on a micromanipulator for precise positioning, and the tip was placed approximately 200 μm away from the target cells, which included HEK293T cells and human retinal ganglion cells (RGCs).

### Statistical Analysis Using Mann-Whitney U Tests with Benjamini-Hochberg False Discovery Rate (FDR) Correction

For comparing peak firing rates in the human retina a comprehensive statistical analysis was performed across different groups (wild-type and Mutant), regions (CP and Macula), and temperatures. For this dataset, the Mann-Whitney U test, a non-parametric method is ideal for comparing two independent samples without assuming a normal distribution of data. Specifically, pairwise comparisons were performed: between CP and Macula regions within each group at each temperature, and between wild-type and Mutant groups within each region and temperature. Recognizing that multiple statistical tests increase the risk of Type I errors (false positives), the code applies the Benjamini-Hochberg False Discovery Rate (FDR) correction to the collection of p-values obtained. By adjusting the p-values, this method balances the need to maintain statistical power—thus not overlooking true positives—while reducing the likelihood of false positives.

### Dynamic Range Calculation, Statistical Analysis, and Phase Plane Analysis

The dynamic range of RGC firing was calculated to assess the responsiveness of cells across different conditions. Peristimulus time histograms (PSTHs) were generated for each recording using a time window of 1.0 s and a bin size of 5 ms. Spike counts were smoothed using a moving average with a window size of five bins to reduce variability and highlight underlying trends. The peak firing rate was identified from these smoothed PSTHs, and the baseline firing rate was calculated as the average firing rate during the last 500 ms of the time window. To account for the logarithmic nature of neuronal response sensitivity, the dynamic range was expressed in decibels (dB) using 20×log10(Peak Firing Rate/Baseline Firing Rate). This approach allowed for normalized comparisons across neurons with varying baseline levels.

For statistical analysis, non-parametric methods were employed due to the non-normal distribution of the dynamic range data. At each temperature, the Kruskal-Wallis test was applied to compare the dynamic ranges among the three groups: wild-type (wild-type), mutant (Δ786-840), and control (Ctrl). To address the issue of multiple comparisons and control the false discovery rate, the Benjamini-Hochberg procedure was applied, as done before, to adjust the p-values obtained from the Kruskal-Wallis tests. For temperatures where significant differences were detected after adjustment (p < 0.05), post-hoc pairwise comparisons were conducted using Tukey’s Honest Significant Difference (HSD) test to identify specific group differences.

To extend the characterization of response dynamics and compare the heat response properties of the channels across groups, phase plane plots were generated. Firing rate data were first z-scored to standardize the responses and facilitate comparison. The time derivatives of the z-scored firing rates were then computed using numerical differentiation (gradient calculation). By plotting the time derivative of the firing rate against the firing rate itself for each group, the phase plane plots illustrated the dynamic evolution of RGC responses to thermal stimuli. Mean trajectories were plotted with shaded error regions representing the standard error of the mean. This analysis is relevant as it provides insights into the kinetics of activation and adaptation of the ion channels in response to heat. Differences in the shape and trajectory of the phase plots among the groups can reveal variations in the temporal dynamic profile of the ion channels.

## Author contributions

M.C. and T.M. performed electrophysiology recordings in human cells; M.C. also conducted recordings in HEK cells and extracted parameters from Boltzmann fits. F.F. carried out rational protein engineering. D.P.M., L.G., and F.K. cultured human retina explants. A.S. coordinated the work on human retina explants from organ donors. G.T.-S. supervised the work, analyzed data, wrote analysis scripts. G.T-S. and B.R., conceived the project. All authors contributed to the manuscript and the supplementary information either through discussions or directly.

---

**Supplementary Table 1.**
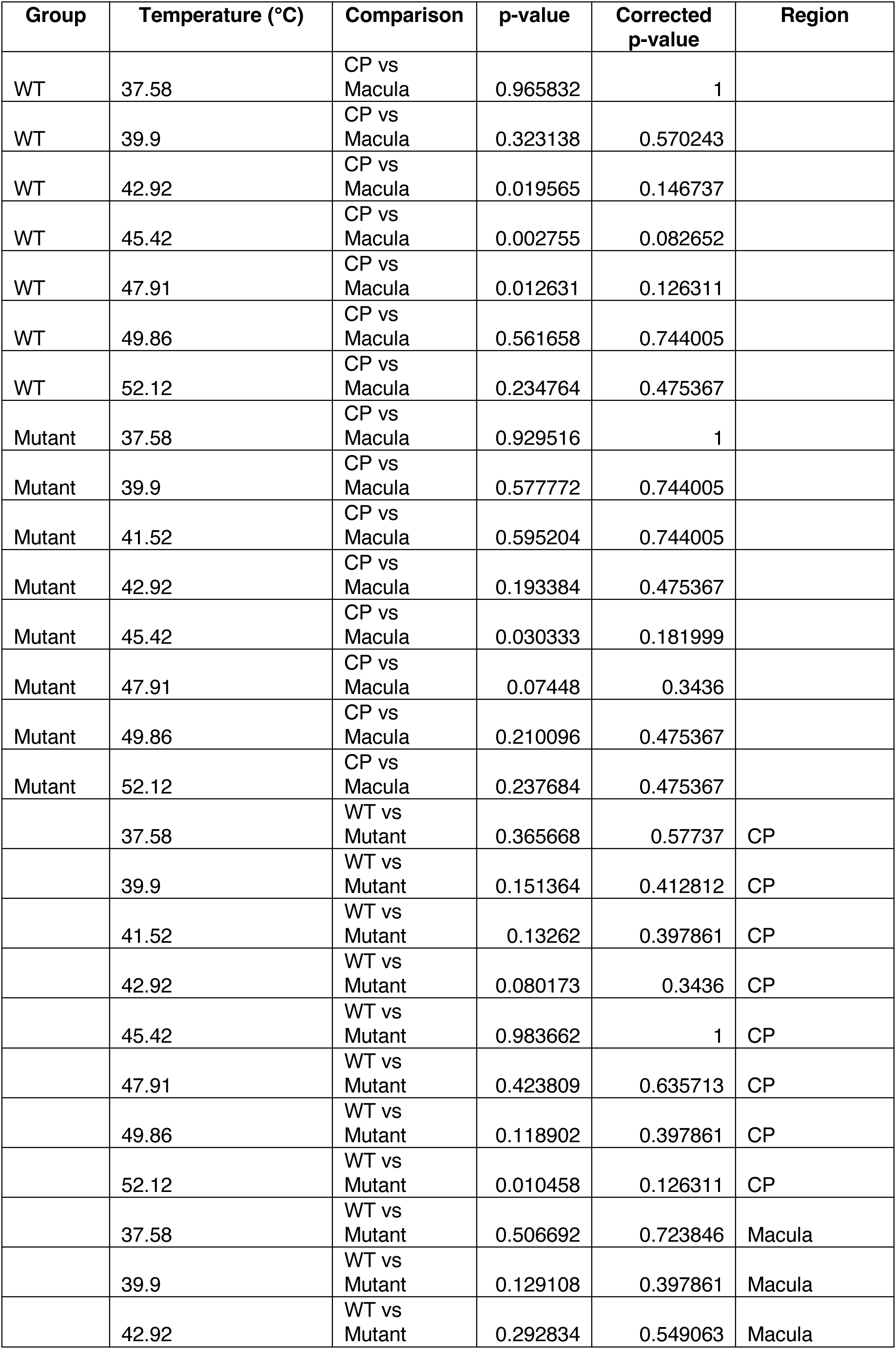
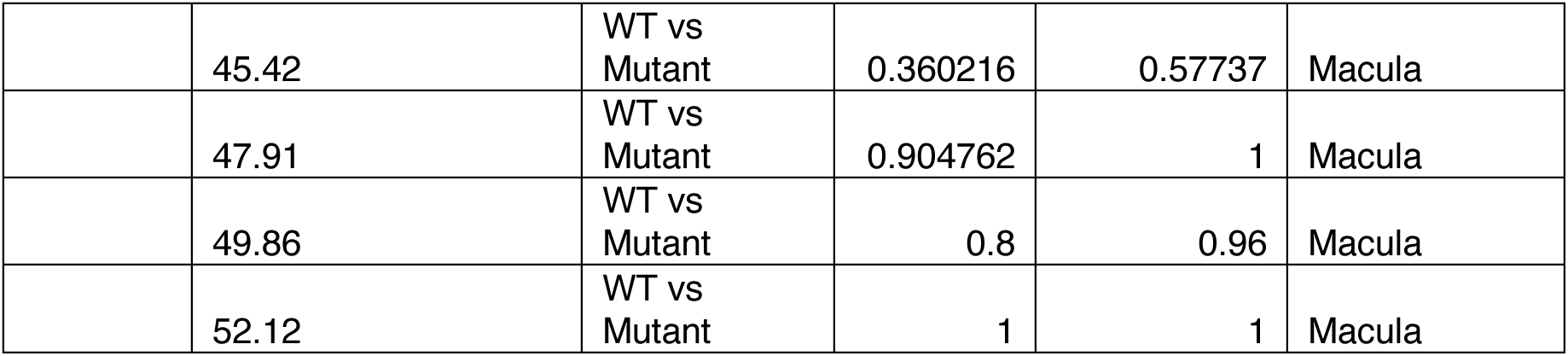
Comparing Peak Firing Rates (Hz) across areas in the human retina and TRPV1 channel phenotype.

| Group | Temperature (°C) | Comparison | p-value | Corrected p-value | Region |
| --- | --- | --- | --- | --- | --- |
| WT | 37.58 | CP vs Macula | 0.965832 | 1 |  |
| WT | 39.9 | CP vs Macula | 0.323138 | 0.570243 |  |
| WT | 42.92 | CP vs Macula | 0.019565 | 0.146737 |  |
| WT | 45.42 | CP vs Macula | 0.002755 | 0.082652 |  |
| WT | 47.91 | CP vs Macula | 0.012631 | 0.126311 |  |
| WT | 49.86 | CP vs Macula | 0.561658 | 0.744005 |  |
| WT | 52.12 | CP vs Macula | 0.234764 | 0.475367 |  |
| Mutant | 37.58 | CP vs Macula | 0.929516 | 1 |  |
| Mutant | 39.9 | CP vs Macula | 0.577772 | 0.744005 |  |
| Mutant | 41.52 | CP vs Macula | 0.595204 | 0.744005 |  |
| Mutant | 42.92 | CP vs Macula | 0.193384 | 0.475367 |  |
| Mutant | 45.42 | CP vs Macula | 0.030333 | 0.181999 |  |
| Mutant | 47.91 | CP vs Macula | 0.07448 | 0.3436 |  |
| Mutant | 49.86 | CP vs Macula | 0.210096 | 0.475367 |  |
| Mutant | 52.12 | CP vs Macula | 0.237684 | 0.475367 |  |
|  | 37.58 | WT vs Mutant | 0.365668 | 0.57737 | CP |
|  | 39.9 | WT vs Mutant | 0.151364 | 0.412812 | CP |
|  | 41.52 | WT vs Mutant | 0.13262 | 0.397861 | CP |
|  | 42.92 | WT vs Mutant | 0.080173 | 0.3436 | CP |
|  | 45.42 | WT vs Mutant | 0.983662 | 1 | CP |
|  | 47.91 | WT vs Mutant | 0.423809 | 0.635713 | CP |
|  | 49.86 | WT vs Mutant | 0.118902 | 0.397861 | CP |
|  | 52.12 | WT vs Mutant | 0.010458 | 0.126311 | CP |
|  | 37.58 | WT vs Mutant | 0.506692 | 0.723846 | Macula |
|  | 39.9 | WT vs Mutant | 0.129108 | 0.397861 | Macula |
|  | 42.92 | WT vs Mutant | 0.292834 | 0.549063 | Macula |
|  | 45.42 | WT vs<br>Mutant | 0.360216 | 0.57737 | Macula |
|  | 47.91 | WT vs<br>Mutant | 0.904762 | 1 | Macula |
|  | 49.86 | WT vs<br>Mutant | 0.8 | 0.96 | Macula |
|  | 52.12 | WT vs<br>Mutant | 1 | 1 | Macula |

**Supplementary Table 2.** Relevant information pertaining to human multi-organ donors.

| **Age** | **Gender** | **Date of donation** | **Transduction**<br>**time** |
| --- | --- | --- | --- |
| 43 | male | 12.06.2023 | 4-6 weeks |
| 53 | male | 14.08.2023 | 4-6 weeks |
| 56 | male | 12.11.2023 | 4-6 weeks |
| 63 | male | 11.06.2024 | 4-6 weeks |
| 63 | female | 30.09.2024 | 4-6 weeks |

## References

1. Busskamp, V., Roska, B. & Sahel, J.-A. Optogenetic Vision Restoration. Cold Spring Harb Perspect Med 14, a041660 (2024).

2. Busskamp, V. et al. Genetic reactivation of cone photoreceptors restores visual responses in retinitis pigmentosa. Science 329, 413–417 (2010).

3. Picaud, S. et al. The primate model for understanding and restoring vision. Proceedings of the National Academy of Sciences of the United States of America 116, 26280 (2019).

4. Sahel, J.-A. et al. Partial recovery of visual function in a blind patient after optogenetic therapy. Nat Med 27, 1223–1229 (2021).

5. Ameln, J. et al. Assessment of local sensitivity in incomplete retinal pigment epithelium and outer retinal atrophy (iRORA) lesions in intermediate age-related macular degeneration (iAMD). BMJ Open Ophth 9, (2024).

6. Bakken, G. S. & Krochmal, A. R. The imaging properties and sensitivity of the facial pits of pitvipers as determined by optical and heat-transfer analysis. Journal of Experimental Biology 210, 2801–2810 (2007).

7. Clark, R. W., Bakken, G. S., Reed, E. J. & Soni, A. Pit viper thermography: the pit organ used by crotaline snakes to detect thermal contrast has poor spatial resolution. Journal of Experimental Biology 225, jeb244478 (2022).

8. Schraft, H. A., Bakken, G. S. & Clark, R. W. Infrared-sensing snakes select ambush orientation based on thermal backgrounds. Sci Rep 9, 3950 (2019).

9. Nelidova, D. et al. Restoring light sensitivity using tunable near-infrared sensors. Science 368, 1108–1113 (2020).

10. Caterina, M. J. et al. The capsaicin receptor: a heat-activated ion channel in the pain pathway. Nature 389, 816–824 (1997).

11. Tominaga, M. et al. The Cloned Capsaicin Receptor Integrates Multiple Pain- Producing Stimuli. Neuron 21, 531–543 (1998).

12. Gracheva, E. O. et al. Molecular basis of infrared detection by snakes. Nature 464, 1006–1011 (2010).

13. Gracheva, E. O. & Bagriantsev, S. N. Evolutionary adaptation to thermosensation. Current Opinion in Neurobiology 34, 67–73 (2015).

14. Gracheva, E. O. et al. Ganglion-specific splicing of TRPV1 underlies infrared sensation in vampire bats. Nature 476, 88–91 (2011).

15. Cao, E., Cordero-Morales, J. F., Liu, B., Qin, F. & Julius, D. TRPV1 channels are intrinsically heat sensitive and negatively regulated by phosphoinositide lipids. Neuron 77, 667–679 (2013).

16. Narasimhan, A. & Jha, K. K. Bio-heat transfer simulation of retinal laser irradiation. International Journal for Numerical Methods in Biomedical Engineering 28, 547–559 (2012).

17. Laursen, W. J., Schneider, E. R., Merriman, D. K., Bagriantsev, S. N. & Gracheva, E. O. Low-cost functional plasticity of TRPV1 supports heat tolerance in squirrels and camels. Proceedings of the National Academy of Sciences 113, 11342–11347 (2016).

18. Neuberger, A. et al. Human TRPV1 structure and inhibition by the analgesic SB- 366791. Nat Commun 14, 2451 (2023).

19. Prescott, E. D. & Julius, D. A Modular PIP2 Binding Site as a Determinant of Capsaicin Receptor Sensitivity. Science 300, 1284–1288 (2003).

20. Yao, J., Liu, B. & Qin, F. Modular thermal sensors in temperature-gated transient receptor potential (TRP) channels. Proceedings of the National Academy of Sciences 108, 11109–11114 (2011).

21. Cowan, C. S. et al. Cell Types of the Human Retina and Its Organoids at Single- Cell Resolution. Cell 182, 1623–1640.e34 (2020).

22. Mirnezami, S. A., Rajaei Jafarabadi, M. & Abrishami, M. Temperature Distribution Simulation of the Human Eye Exposed to Laser Radiation. J Lasers Med Sci 4, 175–181 (2013).

